# iOBPdb – A Database for Experimentally Determined Functional Characterization of Odorant Binding Proteins

**DOI:** 10.1101/2022.06.30.498339

**Authors:** Shalabh Shukla, Oliver Nakano-Baker, Devin MacKenzie, Mehmet Sarikaya

## Abstract

Odorant binding proteins, OBPs, are a diverse family of small, globular, extra-cellular proteins solubilize volatile organic compounds (VOCs) so they can be internalized and transported by an organism. Since their initial discovery in the early eighties ^1^, thousands of OBPs have been identified through genome sequencing and characterized by fluorescence ligand binding assays ^2^. While individual OBPs have been studied in the context of their roles in specific organism, there have been no studies towards the understanding of the comparative structure-function relations of all known OBPs, primarily due to a lack of a centralized database that incorporates the binding affinity with the structure of all OBPs. Incorporating OBP information into a database requires not only an extensive search of all existing resources, but also creating a useful platform that relates sequence structures to target functions. Combining 215 functional studies containing 381 unique OBPs from 91 insect species we created a database, iOBPdb: https://iobpdb.herokuapp.com, of OBP binding affinities for a wide range of VOC targets. We demonstrate here that the construction of this initial database provides powerful search and associative capabilities including interrogating odor binding proteins as clusters and groups by sequence similarity versus protein and target molecular weights, and by the functional groups of the VOC targets. The comparative interrogation of the probe-target recognition allows for a more comprehensive understanding of the underlying structural features of all OBPs that had not been possible by only examining the OBPs individually. We present our results in a variety of phylogenetic representations as well as providing the binding profiles of OBP groups to VOC functional moieties. Potential applications include development of molecular probes for biosensors, novel bioassays and drugs, discovery of novel pesticides which inhibit VOC / OBP interactions, as well providing a foundational basis for the functional understanding of odor sensing and perception in the brain.

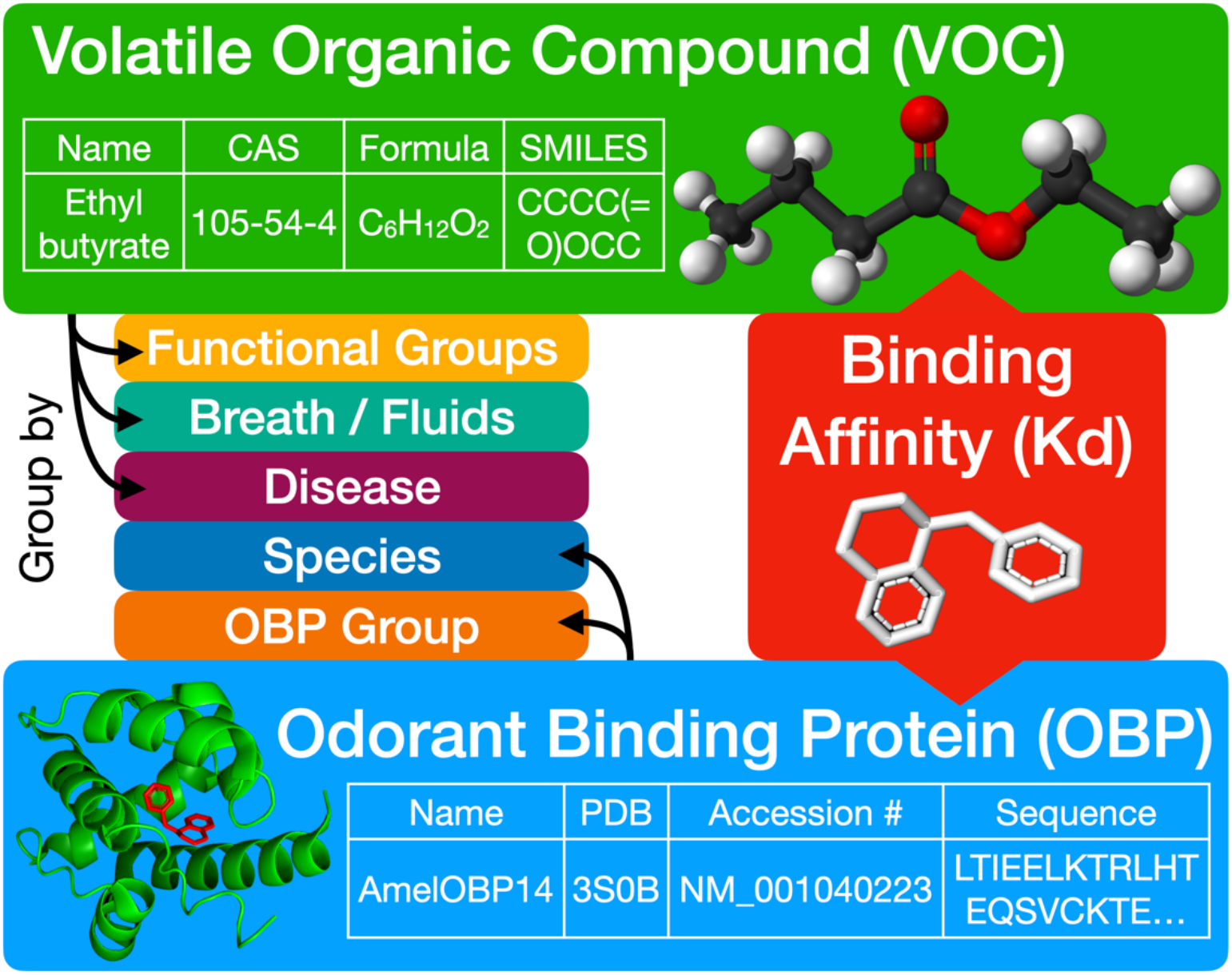

## 1. Introduction and Justification

Odorant binding proteins (OBPs) are a diverse family of small, 10-20 kDa, soluble extracellular target-binding proteins found in both terrestrial vertebrates and invertebrates^2^. Since their initial discovery in insects’ sensillum lymph in 1981 1, thousands of new OBPs have been identified and isolated through genome sequencing and molecular biology approaches. These studies indicate that there is no shared homology between insect and mammalian OBPs. OBPs in mammals are comprised of a beta barrel type structure, whereas the OBPs in insects are a globular structure comprised of alpha helices. Although OBPs are multifaceted in terms of their potential roles in both insects and mammals alike, they are primarily thought to act as odor transporters, solubilizing volatile organic compounds (VOCs) and pheromones from the surrounding air into the aqueous phase of the odor sensory organ, such as the mucus in the nose or sensillum lymph of an antennae^2^. A simplified view of the odorant binding event is shown on figure 1. OBPs transport odorants to sensing neurons which are coated with olfactory receptor (OR) proteins, which can recognize and bind to specific odors, thus signaling an olfactory response^2^. There are also non olfactory related uses of OBPs in insects as OBPs are expressed almost ubiquitously across organs in insects, not just the olfactory organs^3^, and are involved in immunity, mating, moisture detection, signaling molecule transporter, and biochemical inhibitors.

**Figure 1.**
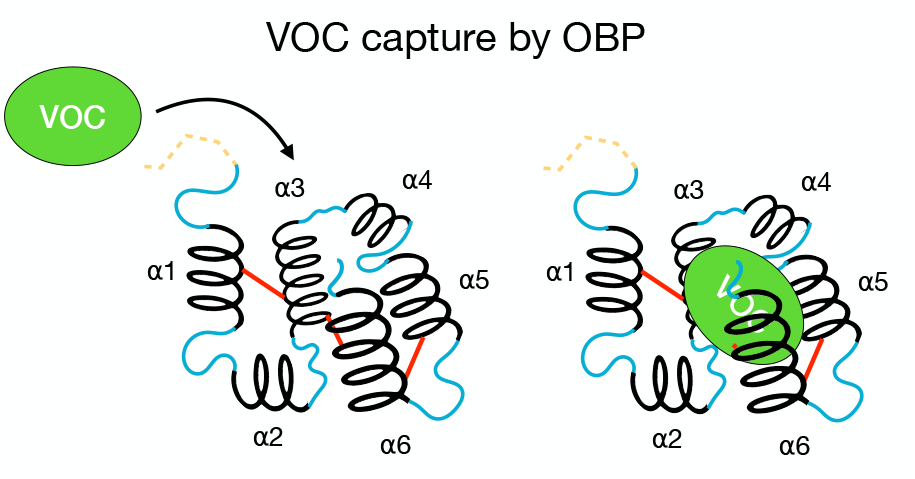
Functional domains of a odor binding protein, OBP, relevant in the capture of a volatile organic compound, VOC.

The literature uses a variety of naming schemes to describe OBP related proteins, such as but not limited to: chemosensory proteins (CSPs), pheromone binding proteins (PBPs), antennal-specific protein (ASPs) antennal binding proteins (ABPXs) and other fringe one off names. Olfactory binding proteins are often sub categorized as either being: classic olfactory binding proteins (classic OBPs), general olfactory binding proteins (GOBPs), atypical olfactory binding proteins (atypical OPBs), minus-C olfactory binding proteins (minus-C OBPs), or plus-C olfactory binding proteins (plus-C OBPs). General odorant binding proteins (GOBPs) are defined based on ubiquitous nature of their expression in both male and female insects. Pheromone binding proteins (PBPs) are a subfamily of OBPs which were identified initially due to their preferential binding to a pheromone in radio-labelled ligand binding studies in the early 1990s^4^. Chemosensory proteins (CSPs) are relatively smaller than other OBPs, share very little sequence homology if any at all, and typically only contain 4 cystines instead of the classic 6 cystines^5^. CSPs are primarily produced in chemosensory organs of insects instead of the antennae^3^.

Inconsistencies of naming arises from the fundamental problem of understanding what parameters makes this large family of thousands of extracellular binding proteins similar or dissimilar from each other. Certain modes of distinguishing these proteins from one another include insect tissue localization, sexual dimorphic expression, insect species of discovery, species of origin of VOC (plant or predator), cystine count in underlying amino acid sequence of binding protein, alpha helices present in folded protein, and preference of binding proteins to certain chemicals or functional groups. While sharing a similar globular structure, insect OBPs differ in terms of conformation, size, and rigidity due to variations in the underlying amino acid sequences, alpha helix count, as well as cystine count (Figure 2). The classic insect OBP contains 6 highly conserved cystine residues which form 3 disulphide bonds. However, there are insect OBP variants with fewer than 5 cystines which only from 2 disulphide bonds and are aptly named minus-C OBPs. Conversely there are insect OBPs with 8 or more cystines which are termed as plus-C OBPs. A special variant of plus-C OBPs are atypical OBPs which typically are 20-30 amino acids longer compared to regular insect OBPs and contain 10 or more cystines. Atypical OBPs are also sometimes referred to as two domain OBPs or double domain OBPs which refers to a fusion protein consisting of two OBPs^6^. The variation in cystine count drastically changes the underlying protein folding, survivability in the extracellular environment and definition of ligand binding pocket. This is not to be confused with dimer OBPs which are two distinct OBP proteins (can be two of the same protein or two different proteins) which pair together in order to sandwich a ligand. The difference is the dimer consists of two distinct proteins whereas the atypical OBP is one large continuous protein.

**Figure 2.**
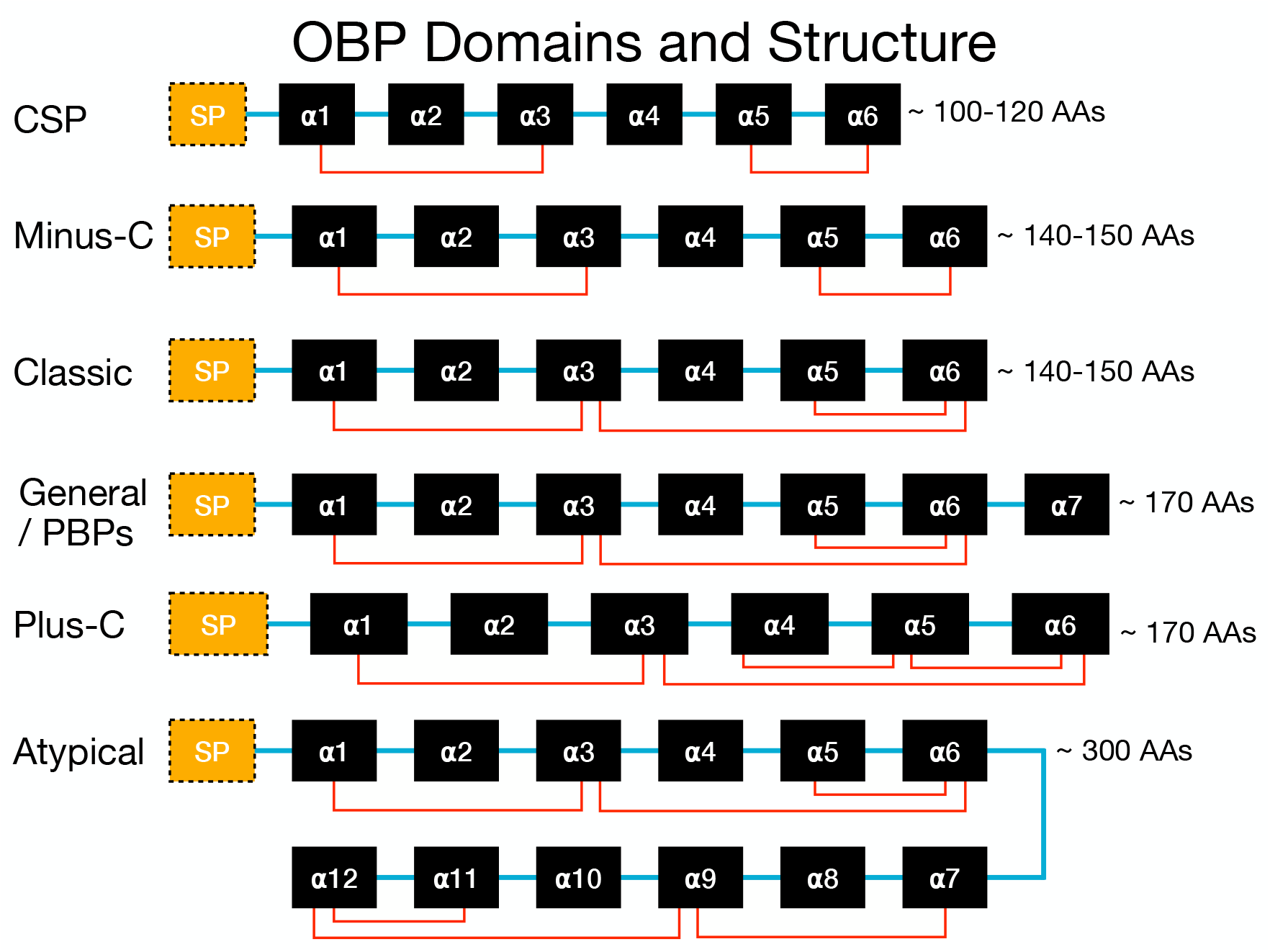
The functional domains of OBPs and their structures. The structural depiction of a classical OBP is with six alpha helices (denoted in black as as α1-α6) and are held in place by three disulphide bonds (denoted in red). The alpha helices are linked by disordered regions (denoted in blue). The signalling peptide (denoted as SP in yellow) at the start of the sequence marks the protein for excretion by the cell and is cleaved before entering the extracellular environment. Different arrangements of OBPs differ in terms of protein size, number of cysteine bonds, and number of alpha helices.

The divergent molecular conformations of insect OBPs alter their function, i.e., the ability to bind to various odors and pheromones. While it is generally agreed that OBPs share specificity for a multitude of different molecules, there are several insect OBPs which exhibit preference for certain molecular moieties and functional groups. Perhaps the most widely known example is LUSH, also known as DmelOBP76a, which was shown to be essential in the recognition of a specific pheromone known as 11-*cis* vaccenyl acetate^7^. When LUSH is knocked out, other OBPs could not compensate in terms of perpetuating signal transduction. The double domain features of atypical OBPs is also speculated to help ensnare signaling molecules in order to prevent signaling action or is potentially needed for binding to unusually large ligands^8^. Conversely fewer cystines appear to be responsible for the relatively broad binding capabilities of CSPs to various ligands.

The specificity of a given OBP to an exclusive target is useful not only in the classification of the OBPs but in terms of their structure-function relationships. The binding of insect OBPs to various molecules have been widely studied using 1-N-phenylnaphthylamine (1-NPN) competitive fluorescent ligand binding assays^9^. 1-NPN is a reporter molecule which provides a continuous baseline signal as it binds to the OBP of interest^10^. Other reporter molecules such as 1-aminoanthracene (1-AMA) are used to study the binding of mammalian OBPs. An odor/pheromone is introduced which competes with 1-NPN in terms of occupation of the ligand binding slot of the OBP causing a decrease in signal. Such fluorescence studies have been accelerating for the last two decades, as the high throughput sequencing of novel insect genomes has made it much easier to identify and isolate novel OBPs.

In this report, we undertake two related studies. First we provide a database from the collected and listed insect OBPs that can be classified in a variety of phylogenic representations. Secondly, we report binding profiles of OBP groups against functional groups found on target VOCs. This should provide a new impetus in functionally classifying OBPs from other organisms and provide a robust platform for odor perception neuroscience, targeted pesticides, and development of biomimetic odorant sensors.

## 2. Materials and Methods

OBP binding data was obtained through an extensive survey of the OBP literature from the last 15 years (see the complete literature sources for the compiled data in the data reporting section). In total, there have been 215 functional studies of 381 unique OBPs from 91 distinct insect species. These functional studies surveyed over 700 different potential binding molecules which are encoded by a CAS (Chemical Abstracts Service) number identifier. Structural information for these molecules were obtained by retrieving the respective molecular formula and SMILES (simplified molecular-input line-entry system) string associated with the CAS identifier through PubChem’s programmatic API, PUG REST^11^. Functional groups present on these molecules were identified by analyzing the underlying SMILE strings. Additional information was obtained from already existing databases and additional literature, making it easier to identify potentially useful OBPs or VOCs which may be pertinent in the realm of human health and biotechnological applications. This includes VOC expression profiles in healthy humans,^12^ VOC profile of various human respiratory diseases,^13–23^ and PDB (Protein Data Bank) entries if fully resolved OBPs exist in RCSB (https://www.rcsb.org), homology matched OBPs exist on the Expasy Swiss-Model database (https://swissmodel.expasy.org), or predicted structure exist in the AlphaFold database (https://alphafold.ebi.ac.uk). For OBPs with no corresponding PDBs in any of the above databases, new PDB structures are denovo generated using AlphaFold V2.0^24^.

To organize information collected from the literature search, a many-to-many relational database was modeled and designed, which we have termed iOBPdb. A static archival version is stored on ResearchWorks, University of Washington’s digital repository for scholarly works. Figure 2 shows the schema for the database. The iOBPdb database is also accessible through a web portal (https://iobpdb.herokuapp.com/). Figure 4 shows how a user can navigate the online database to retrieve information of interest. Figure 4a shows the search page where a user can either search by compound ID or OBP name to retrieve binding data of interest. This binding data can be further filtered by functional groups of VOCs, molecular weight of VOCs, prevalence of VOCs in healthy humans, prevalence of VOCs in disease and many more options. Alternatively, the user can scroll through the compound and OBP browse pages to see the full directory of compounds and OBPs in the database. The search generates a datable of retrieved compounds and OBPs that match the provided descriptions and filters. Clicking on a compound CAS-number (highlighted in blue hypertext format) in the search table navigates the user to a compound information page as seen on Figure 4b. Additionally clicking on an OBP name (highlighted in blue hypertext format) in the search table navigates the user to an OBP information page as seen on Figure 4C. These information pages contain pertinent information such as compound / OBP structure, formula, sequence, binding data etc. Sources for where this information came from is also given on the info pages in doi format.

**Figure 3.**
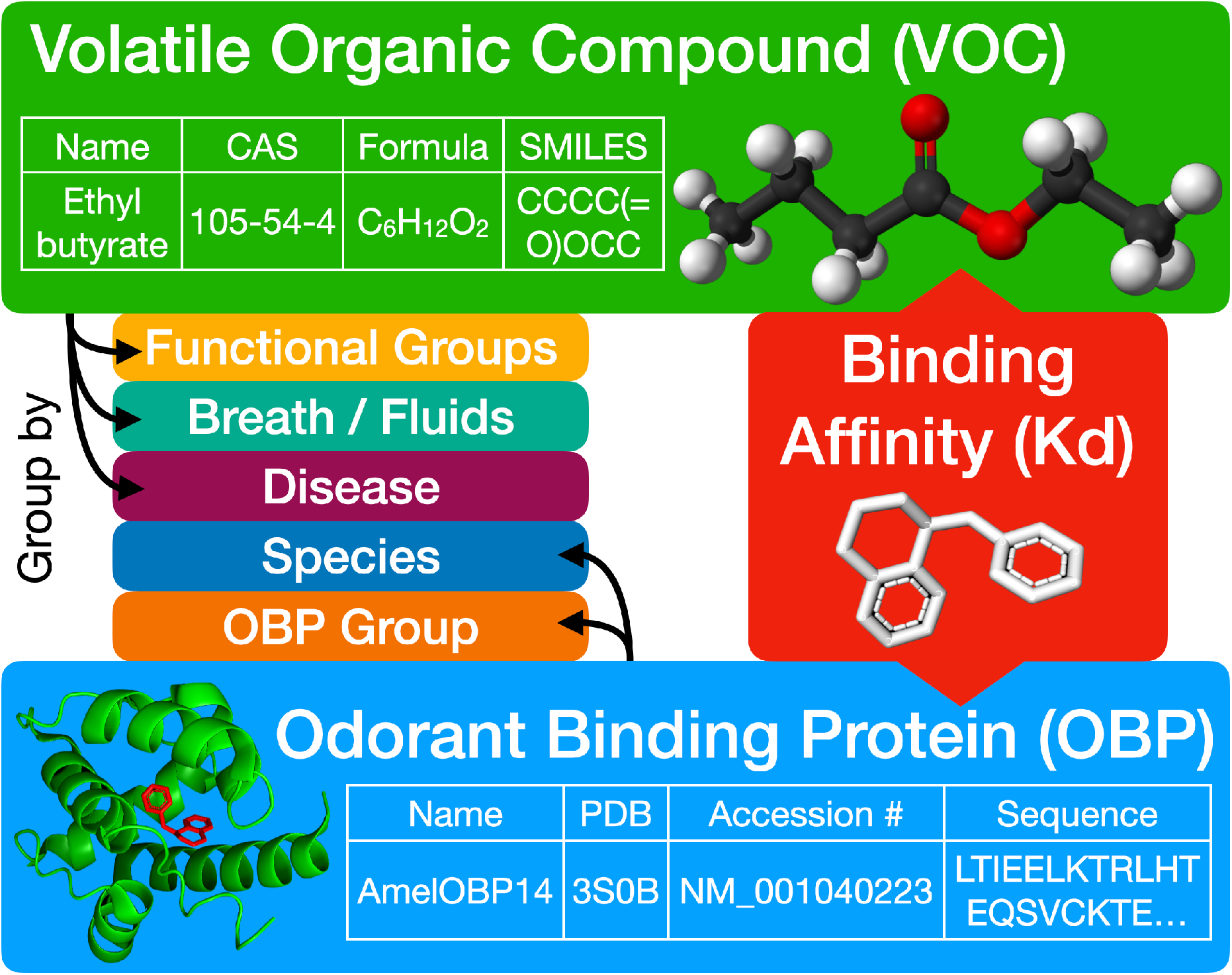
Schematics describing the development of the OBP database for insects, *iOBPdb*. One may search the database by either providing the VOC of interest by providing either a name, CAS number, Molecular Formula, or SMILES string, or alternatively by the OBP of interest by providing either a name, or ascension number. This will return the binding affinity data of OBPs tested against various VOCs per the parameters provided. One can also filter/sort the results by various properties of the VOC such as: its functional groups, presence in healthy humans, as well as presence in diseased humans. These additional filters serve to expedite search for users that may be interested in biotechnological / therapeutic use of OBPs.

**Figure 4.**
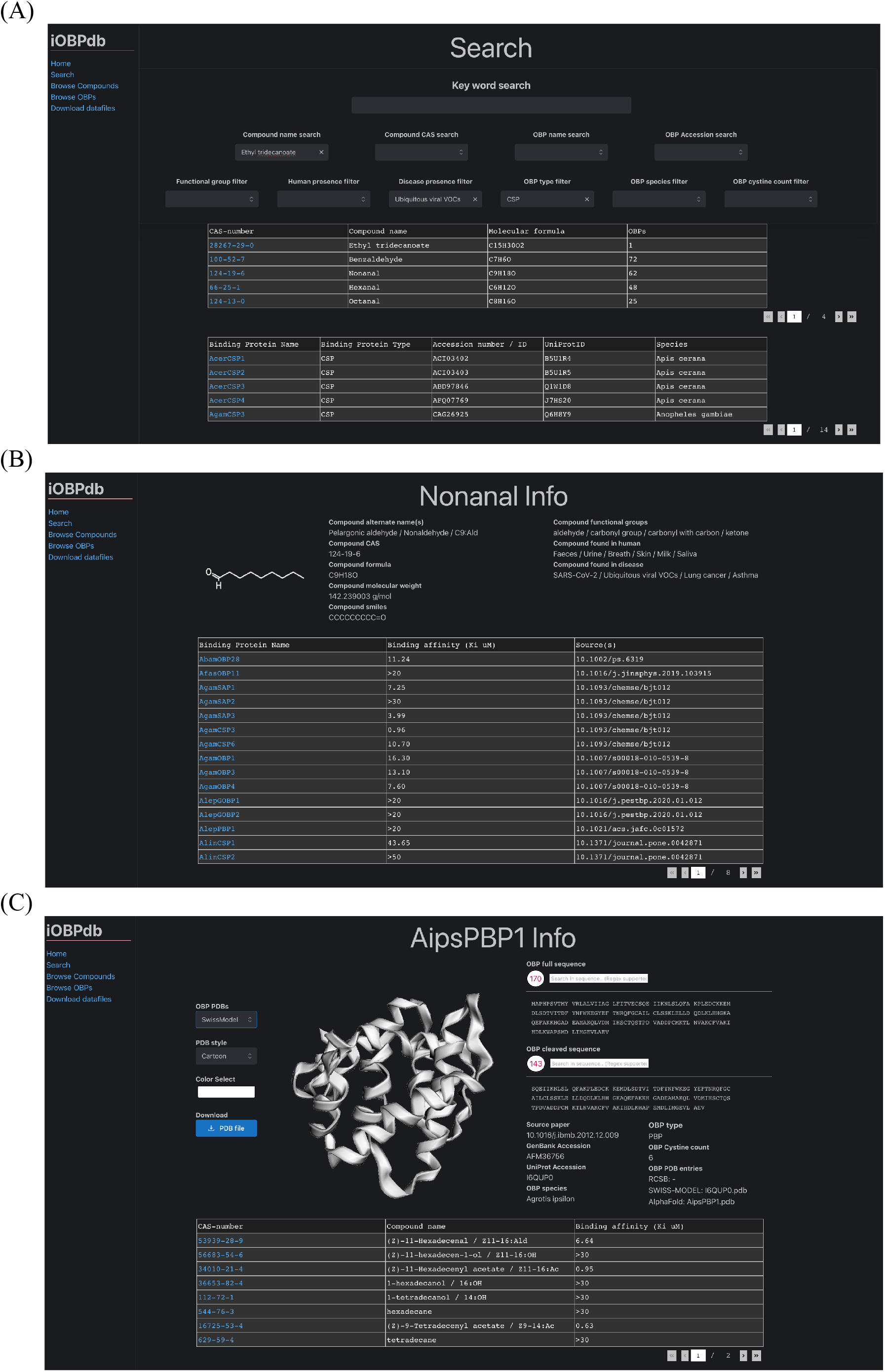
Online iOBPdb web portal (https://iobpdb.herokuapp.com/). Users can search for compounds and OBPs using various key words and filters (4a). Clicking on a compound CAS-number transfers the user to a compound info page (4b). Clicking on an OBP name transfers the user to a OBP info page. For every compound and OBP in the database there is a corresponding information page.

## 3. Results

The acquired OBP information from the literature has been stored in a database called iOBPdb which is intended for use as a robust platform for expansion by including other OBPs, e.g., from mammalians, reptiles, and fish. The database is available for the researchers interested in using the iOBPdb. The comprehensive database is envisioned for use in neuroscience studies via structure-function relationships and the study of odor perception. Additionally the database would also serve as a potent biological resource for biomimetic design of the highly potent molecular probes that would function as the key capture elements in future biosensor architecture.

As examples of exploring the relative similarities of the structure-function relationships among the odor binding proteins from insects, we constructed a phylogenetic map for all OBPs contained in the database in a variety of schemes. For example, the newick tree was generated using Clustal Omega multiple sequence alignment (http://www.ebi.ac.uk/Tools/msa/clustalo/), and the resulting phylogenetic map was constructed and displayed using the iTOL web server. The results of this phylogenetic analysis can be seen in its circular form on Figure 5(a) and unrooted form on Figure 5(b). One can clearly observe that CSPs form a unique clade separate to the minus-C OBPs despite sharing similar cystine counts. Additionally, one can observe two distinct subfamilies of minus-C OBPs, one more closely related to CSPs and the other more closely related to classical OBPs. There is also a distinct sub family of atypical OBPs adjacent to a small sub family of plus-C OBPs. In general, however we observe minimal sequence similarity amongst plus-C OBPs as they are scattered across the phylogenetic tree. This may suggest that they are highly ligand specific as a result. On the other hand, the GOBPs and PBPs form a very distinct monophyletic clade separate form CSPs and OBPs. GOBPs also subdivide into two families termed groups 1 and 2. Classical OBPs have also been subdivided into distinct clades, based on their relative distances, which are designated as clades 1-5.

**Figure 5:**
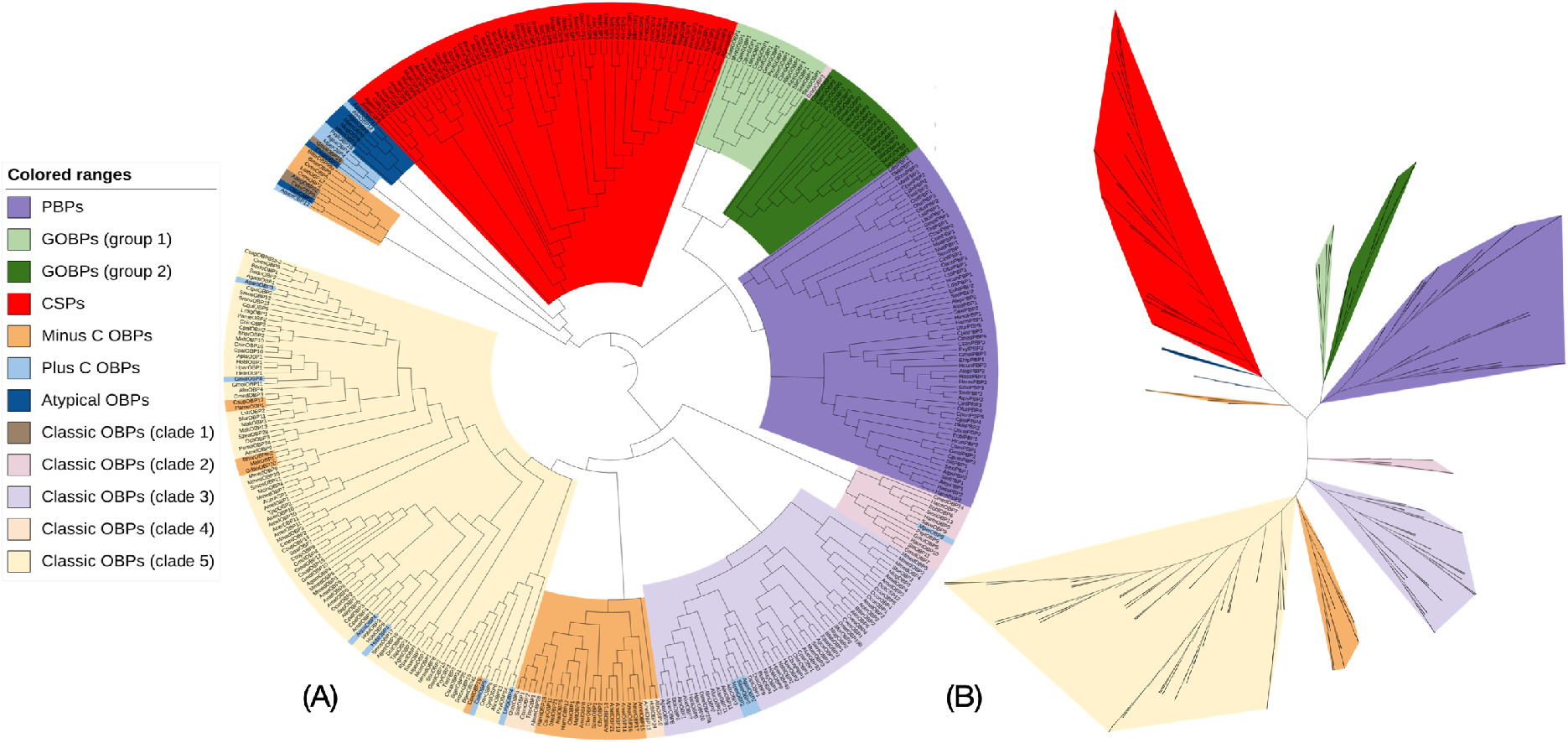
Two types of representations of the insects OBPs: (a) Circular phylogenetic map, and (b) Unrooted phylogenetic map.

The heatmap on Figure 6(a) that overlays the phylogenetic map showcases the distribution of sequence lengths of OBPs. We can clearly observe that CSPs are by far the smallest sub family of proteins, whereas the plus-C and atypical proteins are the largest family. Other interesting features to note are that GOBPs and PBPs are similar in terms of size and are consistently larger than the classical OBPs. Minus-C OBPs and classic OBPs tend to be similar in terms of size.

**Figure 6:**
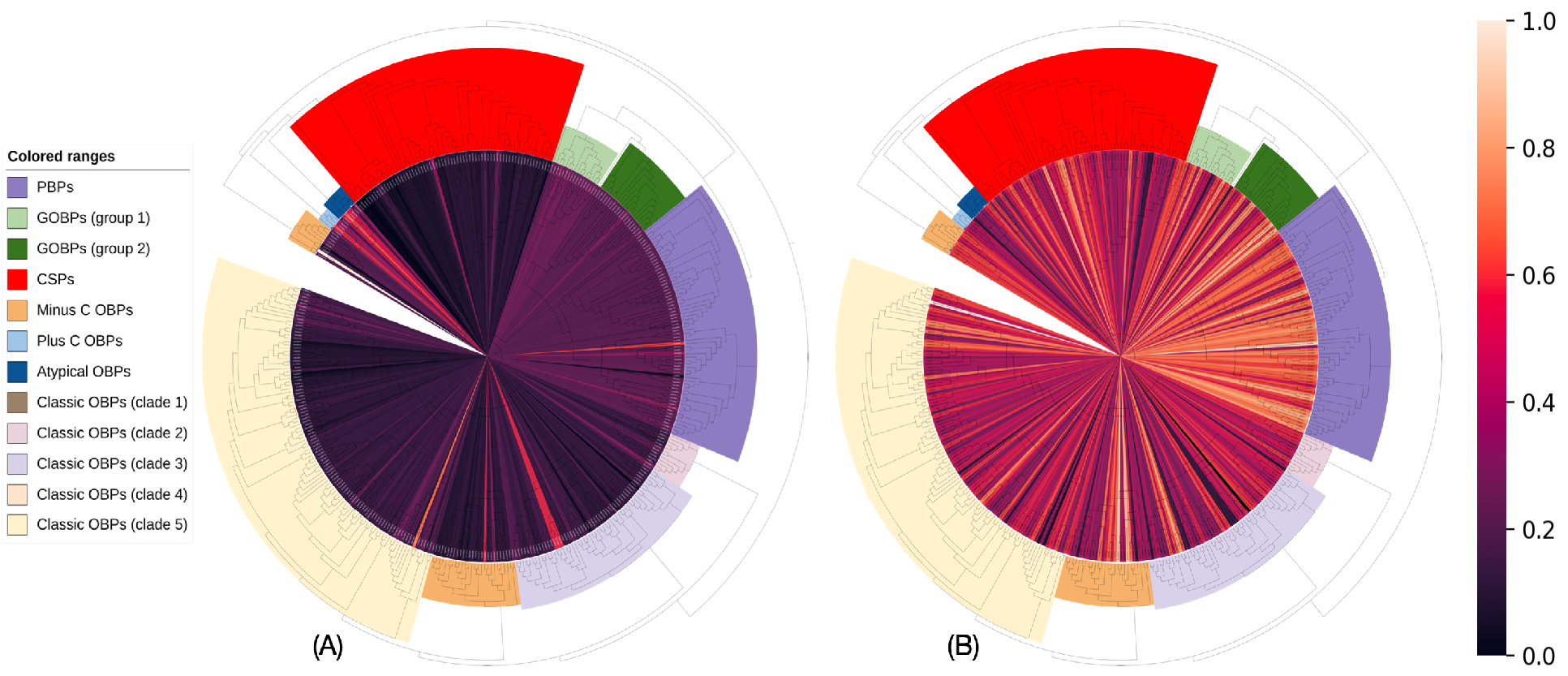
Phylogenetic tree of OBPs generated with respect to the molecular weight (a) Normalized Molecular weight of the OBPs, (b) Normalized VOC Molecular weight preference.

Figure 6(b), a heatmap denoting the preference of each OBP binding to a VOC of a particular molecular weight. This was derived for each OBP by finding the mean molecular weight of all corresponding compounds for where binding data exists. In this case, the PBPs and GOBPs appear to preferentially bind with larger molecular weight VOCs. In the case of PBPs, the binding preference to larger molecular weight VOCs may be an artifact of the experimental bias for testing against larger heavier targets as pheromones tend to be larger molecules. On the other hand, the GOBPs do not have mw-based bias in terms of VOC selection, and they normally tend to preferentially bind to higher molecular weight targets. Classical OBPs, Minus-C OBPs and CSPs do not appear to have a preference in terms of the size of the VOC. Interestingly, Plus-C OBPs seem to display a preference for lower molecular weight targets. This may be a result of the increase in cystines creating more disulfide bonds that restricts the conformation of plus-C OBPs and narrows its binding domain. Similar types of phylogenetic trees could also be generated based on other biochemical or biophysical factors, such as functional groups of VOCs (examples of phylogenetic trees have been produced in the Supplementary Information).

We also present tabulation of mean binding activity of the OBP subfamilies that specifically preferentially bind to certain VOC functional groups (Table 1). Here, all the available VOCs have been considered that had binding activity associated with them. PBPs and GOBPs (group 2 in particular) have an enhanced affinity towards ketones, ethers and aldehydes. Alcohols seem to bind relatively well across several OBP clades, although minus-C OBPs appear to have an enhanced affinity compared to other OBP clades. The CSPs also do not appear to have a strong affinity to any one functional group, which may indicate that the binding activity is broader across the OBPs. An interesting feature that stands out is that phenols preferentially bind to atypical OBPs, Minus – C OBPs and clade 1 OBPs. These are all closely related groups on the phylogenetic map. However the sparsity of representation in some clades, such as clade 1 OBPs and atypical OBPs, makes coming to any definitive conclusions on clade-functional group specificity incredibly spurious. This analysis would greatly benefit from additional OBP functional studies. Further tables of OBP group binding profiles for VOC specific targets (i.e. benzaldehyde, nonanal etc.) can be found in the Supplementary Information.

**Table 1:**
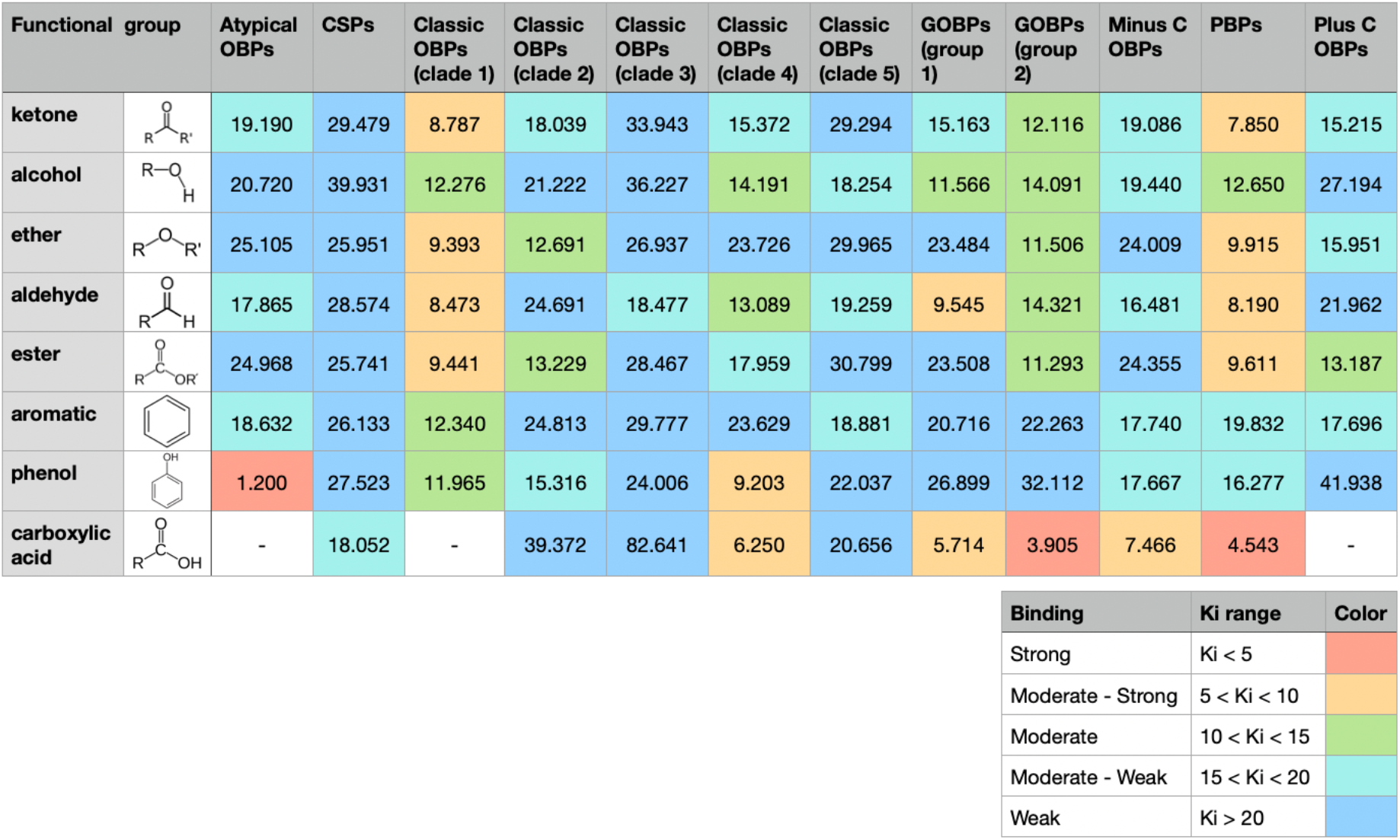
Mean binding affinity (Ki) of functional moieties by OBP groups

| Functional group |  | Atypical OBPs | CSPs | Classic OBPs (clade 1) | Classic OBPs (clade 2) | Classic OBPs (clade 3) | Classic OBPs (clade 4) | Classic OBPs (clade 5) | GOBPs (group 1) | GOBPs (group 2) | Minus C OBPs | PBPs | Plus C OBPs |
| --- | --- | --- | --- | --- | --- | --- | --- | --- | --- | --- | --- | --- | --- |
| ketone |  | 19.190 | 29.479 | 8.787 | 18.039 | 33.943 | 15.372 | 29.294 | 15.163 | 12.116 | 19.086 | 7.850 | 15.215 |
| alcohol |  | 20.720 | 39.931 | 12.276 | 21.222 | 36.227 | 14.191 | 18.254 | 11.566 | 14.091 | 19.440 | 12.650 | 27.194 |
| ether |  | 25.105 | 25.951 | 9.393 | 12.691 | 26.937 | 23.726 | 29.965 | 23.484 | 11.506 | 24.009 | 9.915 | 15.951 |
| aldehyde |  | 17.865 | 28.574 | 8.473 | 24.691 | 18.477 | 13.089 | 19.259 | 9.545 | 14.321 | 16.481 | 8.190 | 21.962 |
| ester |  | 24.968 | 25.741 | 9.441 | 13.229 | 28.467 | 17.959 | 30.799 | 23.508 | 11.293 | 24.355 | 9.611 | 13.187 |
| aromatic |  | 18.632 | 26.133 | 12.340 | 24.813 | 29.777 | 23.629 | 18.881 | 20.716 | 22.263 | 17.740 | 19.832 | 17.696 |
| phenol |  | 1.200 | 27.523 | 11.965 | 15.316 | 24.006 | 9.203 | 22.037 | 26.899 | 32.112 | 17.667 | 16.277 | 41.938 |
| carboxylic acid |  | - | 18.052 | - | 39.372 | 82.641 | 6.250 | 20.656 | 5.714 | 3.905 | 7.466 | 4.543 | - |

## 4. Discussions

In the results, we have presented the outcome of the OBP-VOC based ligand-receptor interactions and presented them in a variety of phylogenetic visualization as well as provided the database algorithm that facilitates the useful classification of these unique molecular interactions. The expectation here is that the initial version of iOBPdb would trigger, as a working platform for further studies in the classification of the odor binding proteins in other organism, such as mammalians, reptiles, and fishes.

It has been well known that the organismal olfactory and gustatory systems can detect and discriminate volatile and solubilized molecules at low concentrations in air and in water for smell and taste, respectively. As schematically simplified in Figures 1 and 2, the process of target detection involves capturing of the target molecule that triggers a biological sensing mechanism by which a signal is transferred to and is recognized by the brain that could discriminate even minute differences in the elemental compositions and structures of thousands of molecules all at once. Gaining insights into the detailed understanding of the processes of sensing have far reaching implications. Firstly, molecular detection has significance in the fundamental processes that underly the mechanisms of molecular recognition and signal transduction processes, e.g., in olfactory systems, specialized by a variety of classes of organisms. Secondly, a database of OBP / VOC profiles for a variety of insect species can facilitate the creation and design of pesticides which can leverage similarities in binding profiles of certain VOC functional group moieties to create OBP specific inhibitors. Finally, the iOBPdb database could unlock the potential use of OBPs as molecular biomimetic lessons for the design of engineered molecular probes.

Despite numerous functional binding studies, relatively few applications of insect OBPs in biosensors were reported^25^. OBPs are small globular proteins are robust enough to stand up to wide ranges of pH and temperatures (even to 80–100°C)^26^, without denaturing and losing their binding properties. In addition, OBPs are easier to isolate and purify in the process of production compared with membrane protein ORs, because ORs are G protein-coupled receptors, and have seven transmembrane α helix^27^. All of these will greatly enhance the practicability of those materials using in sensors. Therefore, OBPs are excellent candidates as biological elements in the development of olfactory-derived biosensors.

## 5. Conclusions and Future Prospects

In insects, the recognition of odor is accomplished by odor binding proteins in binding to and carrying the specific odor to olfactory membrane-bound proteins in the antenna, the first step in odor perception. Although there are considerable studies in odor recognition during the last two decades there is a lack of comprehensive database of OBP-odor interactions. iOBPdb provides a novel platform for broad study of odorant binding proteins and volatile organic compounds interactions. The iOBPdb:

– Provides a comprehensive database, with compiled OBPs with binding specificity to VOC, in insects, which had not been possible before
– Provides a platform to compare multiple OBPs from different insect species, allowing robust comparative analysis

We hope that this study will accelerate the understanding of the physiochemical and neurological basis for odor perception in the brain, discovery of novel targeted pesticides for specific insect species, and applications for OBPs and molecular probes derived from OBPs in devices such as biosensors for volatile organic compound identification in diseased or toxin affected individuals, odor specific analytical probes, and many more possibilities enabled by the evolved, rich functionality of olfactory systems.

## 6. Acknowledgements

We thank Igor, Sami, Tatum, Sid, Le, Richard, Hanson. Partial funding for this work was received from the WE-REACH program at the University of Washington that is supported through the RADx RAD program at NIDCR/NIH.

## 7. Data Sharing

The archival CSV file containing all of the data stored on iOBPdb is available here: https://depts.washington.edu/dennig2/phpmyadmin/

iOBPdb can be accessed online here: https://iobpdb.herokuapp.com/

