## Supplemental figures for "iOBPdb – A Database for Experimentally Determined Functional Characterization of Odorant Binding Proteins"

### Distribution of VOC molecular weights tested for by OBP category

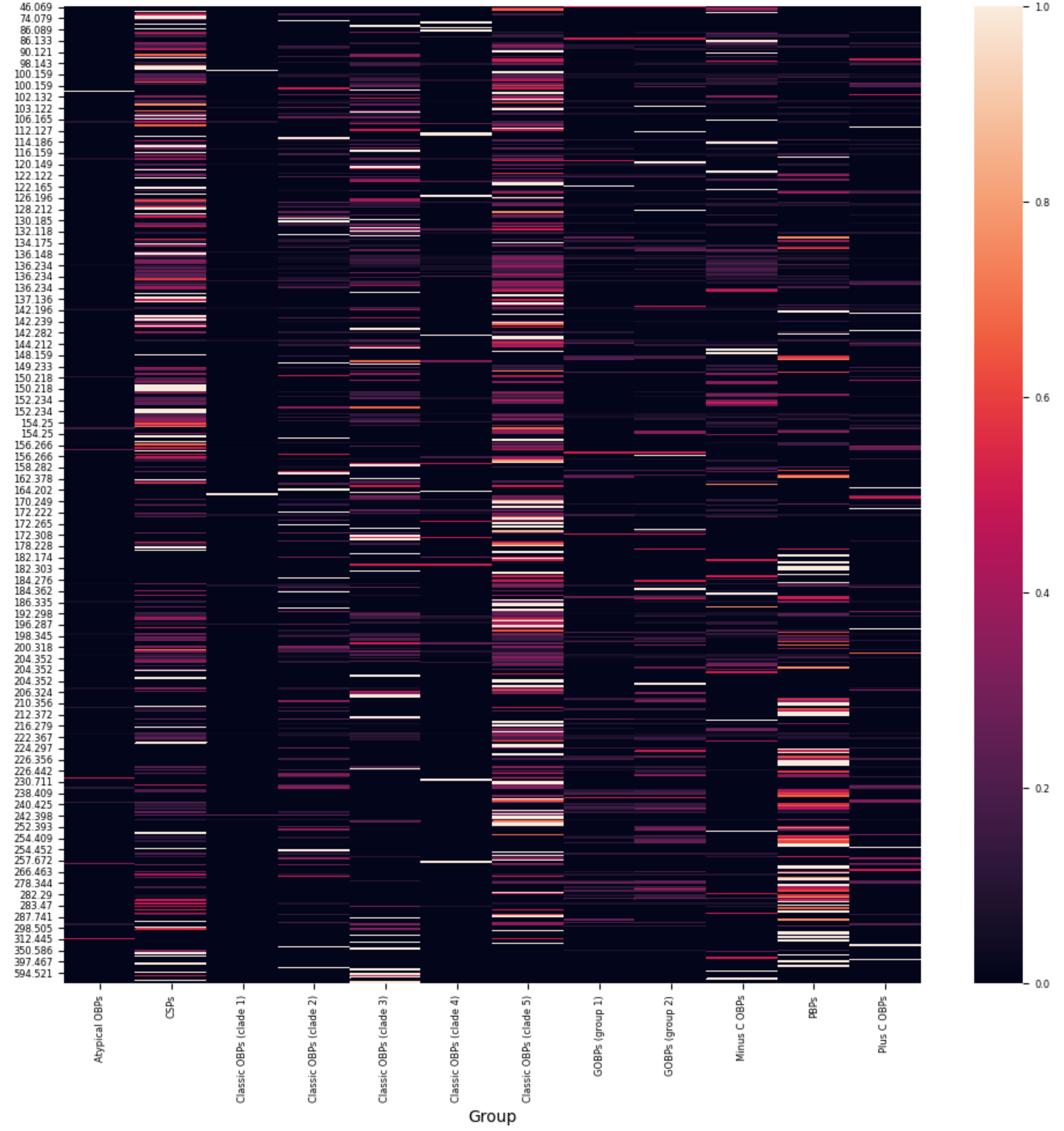

total VOCs tested against by functional group

| Group | Total VOCs tested against | ketone | alcohol | ether | aldehyde | ester | aromatic | phenol | carboxylic acid |
|---|---|---|---|---|---|---|---|---|---|
| **Atypical OBPs** | **184** | 16 | 9 | 6 | 5 | 5 | 8 | 1 | 0 |
| **CSPs** | **1462** | 398 | 254 | 159 | 155 | 133 | 210 | 46 | 18 |
| **Classic OBPs (clade 1)** | **43** | 17 | 8 | 10 | 6 | 9 | 7 | 2 | 0 |
| **Classic OBPs (clade 2)** | **287** | 114 | 57 | 58 | 43 | 49 | 59 | 6 | 5 |
| **Classic OBPs (clade 3)** | **960** | 225 | 136 | 102 | 65 | 85 | 109 | 42 | 16 |
| **Classic OBPs (clade 4)** | **82** | 26 | 10 | 13 | 9 | 10 | 12 | 3 | 2 |
| **Classic OBPs (clade 5)** | **2083** | 483 | 255 | 257 | 124 | 229 | 175 | 40 | 15 |
| **GOBPs (group 1)** | **303** | 92 | 52 | 33 | 39 | 32 | 31 | 5 | 5 |
| **GOBPs (group 2)** | **372** | 135 | 77 | 56 | 49 | 55 | 32 | 6 | 11 |
| **Minus C OBPs** | **493** | 123 | 68 | 48 | 41 | 35 | 33 | 12 | 7 |
| **PBPs** | **821** | 352 | 184 | 154 | 134 | 140 | 73 | 15 | 33 |
| **Plus C OBPs** | **344** | 92 | 47 | 48 | 30 | 44 | 41 | 6 | 0 |

percentage of functional group present in VOCs tested

| Group | ketone | alcohol | ether | aldehyde | ester | aromatic | phenol | carboxylic acid |
|---|---|---|---|---|---|---|---|---|
| **Atypical OBPs** | 8.696% | 4.891% | 3.261% | 2.717% | 2.717% | 4.348% | 0.543% | 0.000% |
| **CSPs** | 27.223% | 17.373% | 10.876% | 10.602% | 9.097% | 14.364% | 3.146% | 1.231% |
| **Classic OBPs (clade 1)** | 39.535% | 18.605% | 23.256% | 13.953% | 20.930% | 16.279% | 4.651% | 0.000% |
| **Classic OBPs (clade 2)** | 39.721% | 19.861% | 20.209% | 14.983% | 17.073% | 20.557% | 2.091% | 1.742% |
| **Classic OBPs (clade 3)** | 23.438% | 14.167% | 10.625% | 6.771% | 8.854% | 11.354% | 4.375% | 1.667% |
| **Classic OBPs (clade 4)** | 31.707% | 12.195% | 15.854% | 10.976% | 12.195% | 14.634% | 3.659% | 2.439% |
| **Classic OBPs (clade 5)** | 23.188% | 12.242% | 12.338% | 5.953% | 10.994% | 8.401% | 1.920% | 0.720% |
| **GOBPs (group 1)** | 30.363% | 17.162% | 10.891% | 12.871% | 10.561% | 10.231% | 1.650% | 1.650% |
| **GOBPs (group 2)** | 36.290% | 20.699% | 15.054% | 13.172% | 14.785% | 8.602% | 1.613% | 2.957% |
| **Minus C OBPs** | 24.949% | 13.793% | 9.736% | 8.316% | 7.099% | 6.694% | 2.434% | 1.420% |
| **PBPs** | 42.875% | 22.412% | 18.758% | 16.322% | 17.052% | 8.892% | 1.827% | 4.019% |
| **Plus C OBPs** | 26.744% | 13.663% | 13.953% | 8.721% | 12.791% | 11.919% | 1.744% | 0.000% |

Mean Binding Affinity (Ki) of VOCs by OBP (only for VOCs with binding affinity < 20 Ki)-1

| VOC | | Atypical OBPs | CSPs | Classic OBPs (clade 1) | Classic OBPs (clade 2) | Classic OBPs (clade 3) | Classic OBPs (clade 4) | Classic OBPs (clade 5) | GOBPs (group 1) | GOBPs (group 2) | Minus C OBPs | PBPs | Plus C OBPs |
|---|---|---|---|---|---|---|---|---|---|---|---|---|---|
| beta ionone | 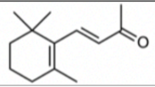 | 13.170 | 10.202 | - | 13.010 | 9.054 | 23.800 | 13.547 | 14.003 | 8.933 | 9.951 | 9.924 | 19.750 |
| Z-3-hexen-1-ol | 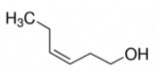 | 39.500 | 23.458 | 24.580 | 11.833 | 17.939 | - | 27.982 | 16.415 | 11.595 | 20.517 | 10.377 | 16.525 |
| benzaldehyde | 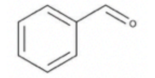 | - | 56.136 | 23.840 | 52.788 | 22.887 | - | 23.735 | 15.876 | 40.670 | 13.363 | 16.220 | 12.003 |
| beta myrcene | 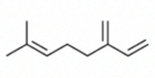 | - | 20.778 | - | 12.580 | 18.134 | 14.525 | 17.211 | 25.827 | 24.946 | 13.745 | 10.750 | 9.700 |
| (+/-) nerolidol | 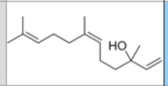 | 28.825 | 12.411 | 25.670 | 15.433 | 16.561 | 5.890 | 13.224 | - | 6.875 | 15.970 | 8.884 | 15.560 |
| (E)-2-Hexenal | 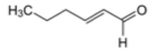 | - | 25.066 | 20.930 | 23.430 | 24.092 | 20.500 | 24.053 | 18.630 | 20.913 | 27.714 | 22.441 | 15.185 |
| Cis-3-Hexenylacetate | 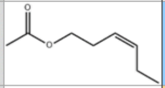 | 34.500 | 14.075 | 15.335 | 20.050 | 28.344 | - | 19.413 | 19.717 | 12.308 | 17.823 | 26.324 | 16.657 |
| nonanal | 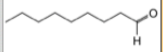 | 6.6 | 22.737 | 4.6 | 24.863 | 16.372 | 11.240 | 22.088 | 30.055 | 45.517 | 13.510 | 8.257 | 16.18 |
| DL linalool | 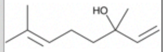 | - | 24.030 | - | 35.532 | 18.453 | 12.650 | 16.451 | 25.365 | 14.365 | 10.428 | 13.687 | 21.150 |
| methylsalicylate | 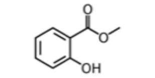 | - | 27.828 | 20.250 | 10.865 | 26.863 | - | 32.289 | 45.770 | 42.930 | 48.100 | 18.525 | 18.790 |

Mean Binding Affinity (Ki) of VOCs by OBP (only for VOCs with binding affinity < 20 Ki)

| VOC | | Atypical OBPs | CSPs | Classic OBPs (clade 1) | Classic OBPs (clade 2) | Classic OBPs (clade 3) | Classic OBPs (clade 4) | Classic OBPs (clade 5) | GOBPs (group 1) | GOBPs (group 2) | Minus C OBPs | PBPs | Plus C OBPs |
|---|---|---|---|---|---|---|---|---|---|---|---|---|---|
| beta ionone | 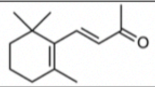 | 13.170 | 7.849 | - | 5.320 | 7.632 | - | 10.159 | 14.003 | 8.933 | 9.951 | 8.901 | 7.090 |
| Z-3-hexen-1-ol | 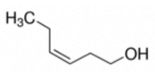 | - | 14.095 | - | 11.833 | 11.094 | - | 14.715 | 11.025 | 9.672 | 6.375 | 10.377 | 11.767 |
| benzaldehyde | 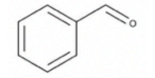 | - | 10.788 | - | 11.135 | 12.347 | - | 11.527 | 11.805 | 11.200 | 2.645 | 8.427 | 7.580 |
| beta myrcene | 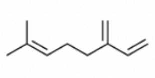 | - | 9.295 | - | 12.580 | 10.333 | 14.525 | 12.223 | 11.680 | 10.753 | 8.237 | 8.258 | 9.700 |
| (+/-) nerolidol | 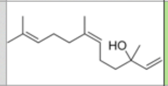 | 14.950 | 8.982 | - | 11.380 | 10.858 | 5.890 | 7.697 | - | 6.875 | 3.390 | 8.884 | 12.140 |
| (E)-2-Hexenal | 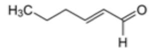 | - | 10.577 | - | 15.055 | 10.053 | 13.690 | 10.338 | 16.060 | 6.200 | 8.235 | 9.910 | 10.928 |
| Cis-3-Hexenylacetate | 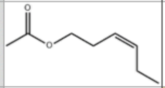 | - | 12.277 | 5.000 | 13.690 | 15.168 | - | 11.173 | 15.490 | 12.308 | 4.085 | 8.178 | 16.657 |
| nonanal | 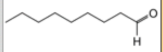 | 6.6 | 7.245 | 4.6 | 19.590 | 9.993 | 11.240 | 10.826 | 5.690 | - | 5.710 | 8.257 | 16.18 |
| DL linalool | 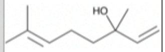 | - | 11.524 | - | 12.025 | 9.350 | 5.000 | 8.720 | 6.700 | 11.477 | 5.530 | 10.885 | 18.300 |
| methylsalicylate | 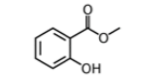 | - | 12.304 | - | 10.865 | 11.645 | - | 12.771 | - | 7.970 | - | 7.550 | 18.790 |

| Binding | Ki range | Color |
|---|---|---|
| Strong | Ki < 5 | |
| Moderate - Strong | 5 < Ki < 10 | |
| Moderate | 10 < Ki < 15 | |
| Moderate - Weak | 15 < Ki < 20 | |
| Weak | Ki > 20 | |

### Refrence Phlyogenetic Map of OBPs

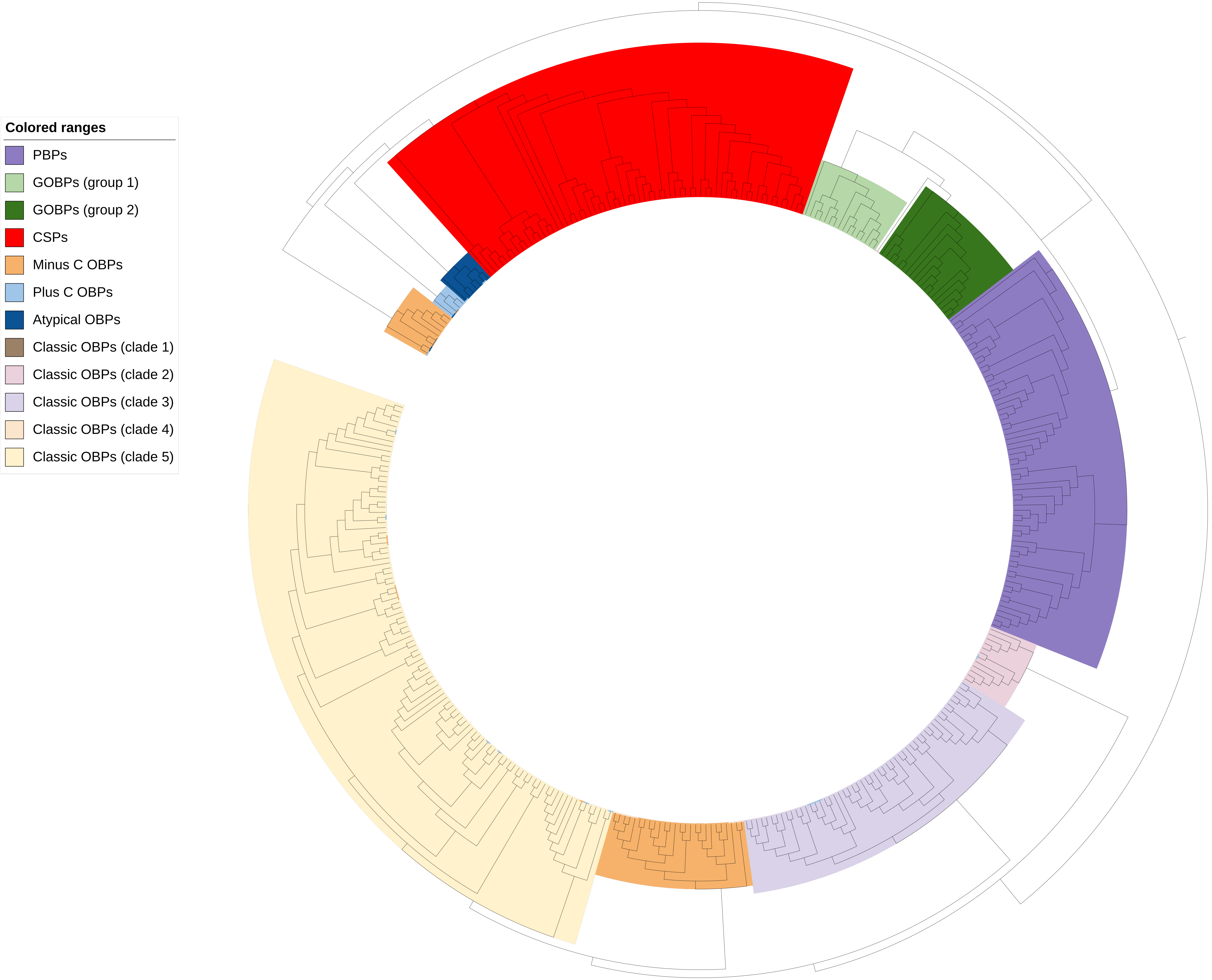

### Ketone preference

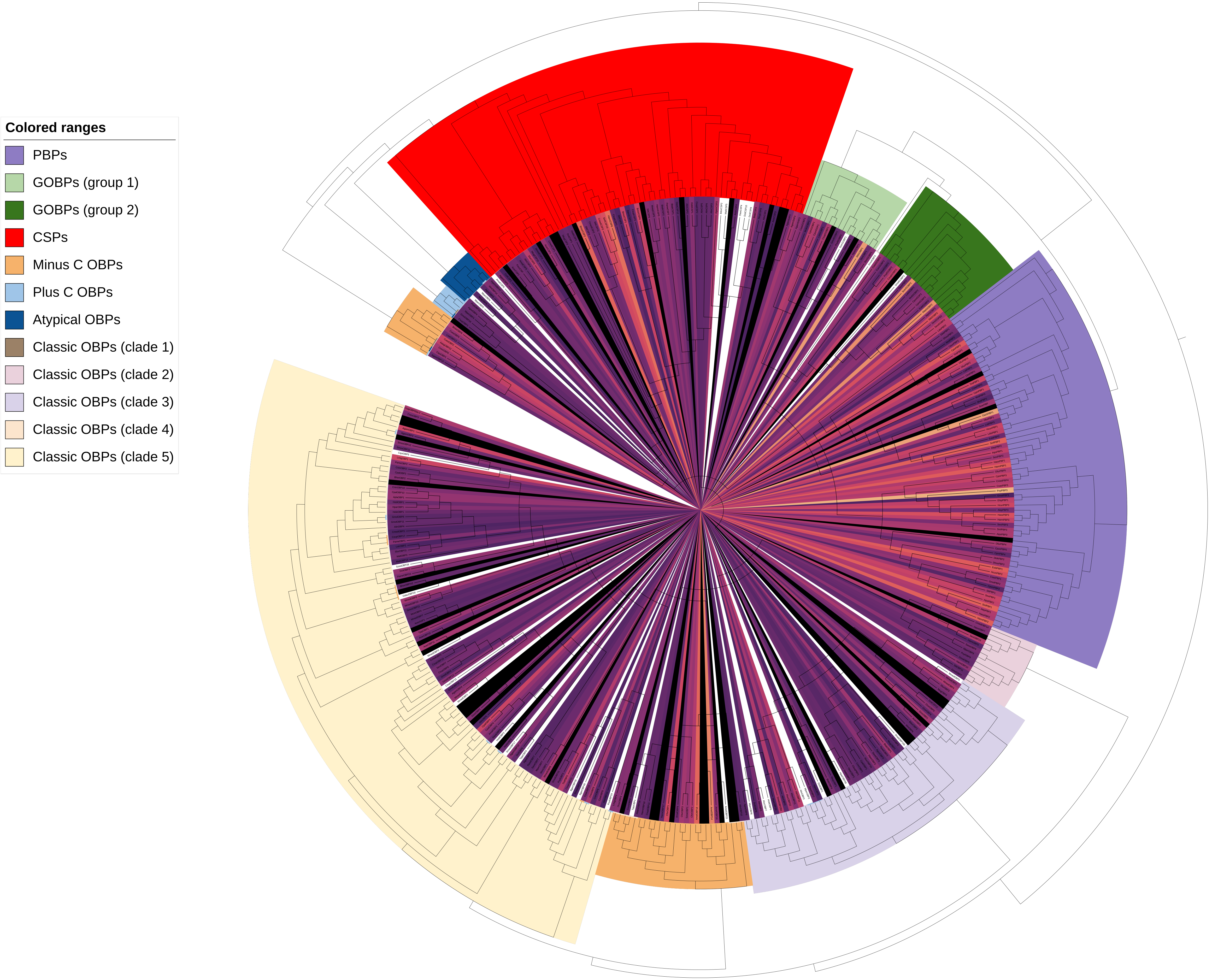

### Alcohol preference

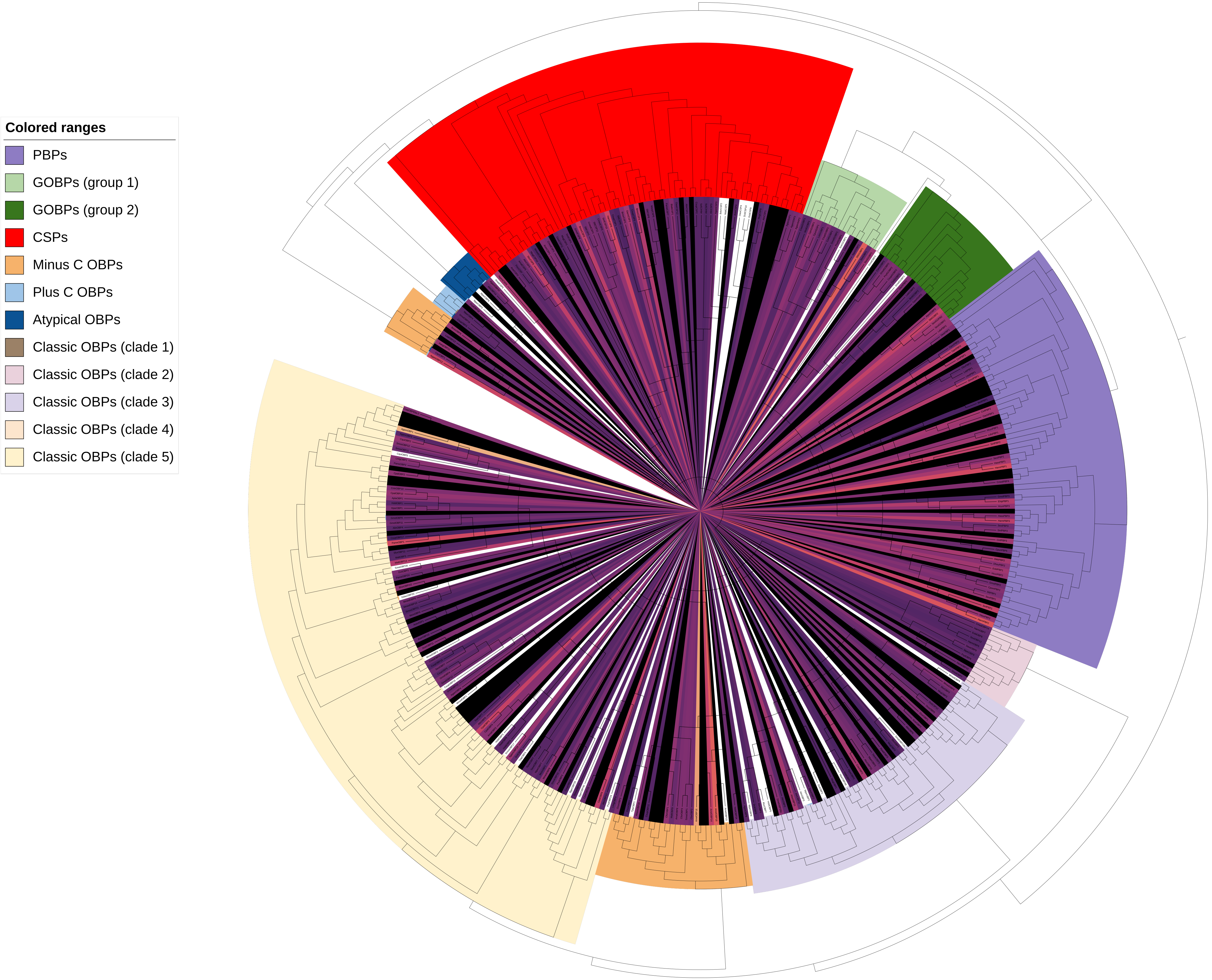

### Ether preference

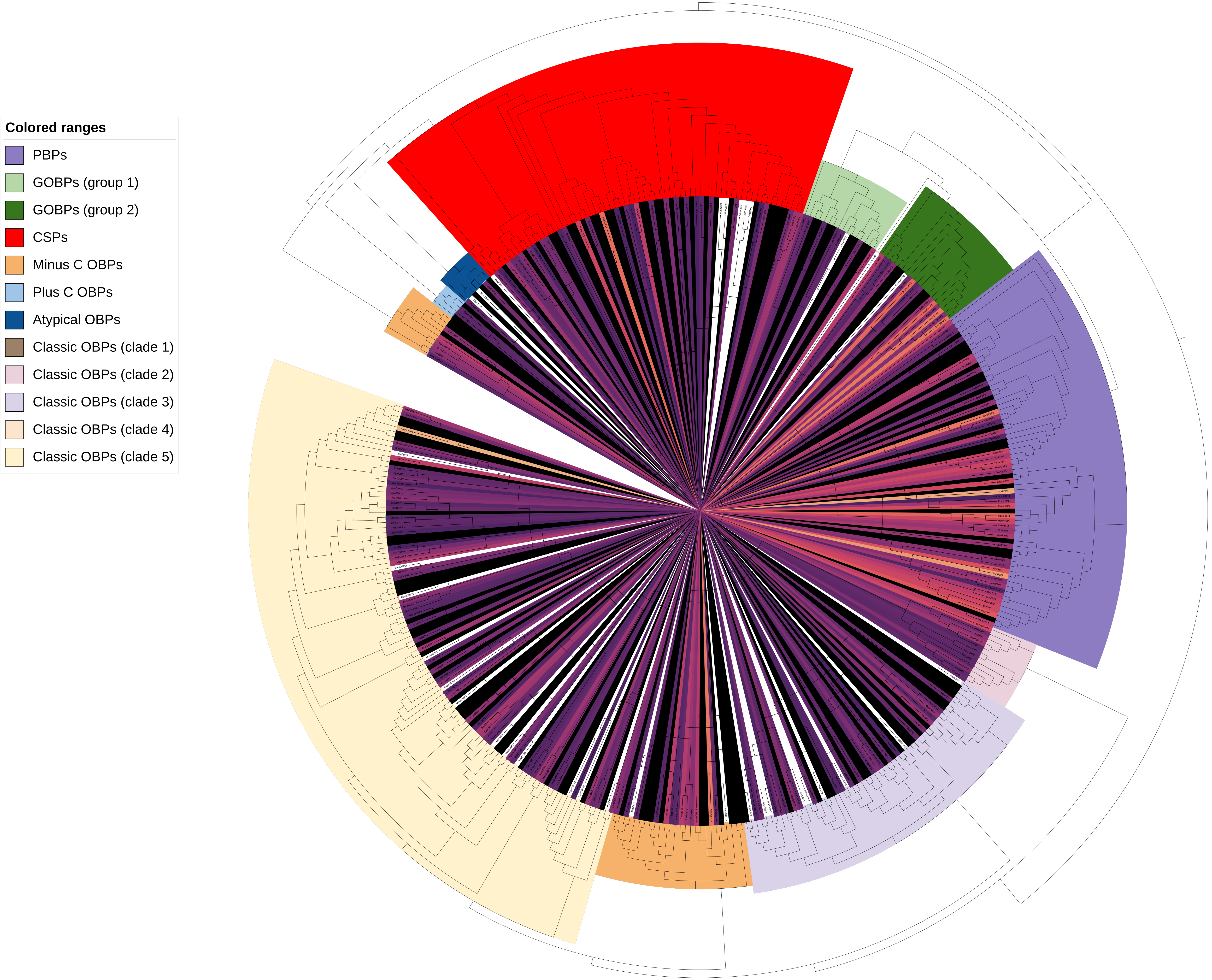

### Aldehyde preference

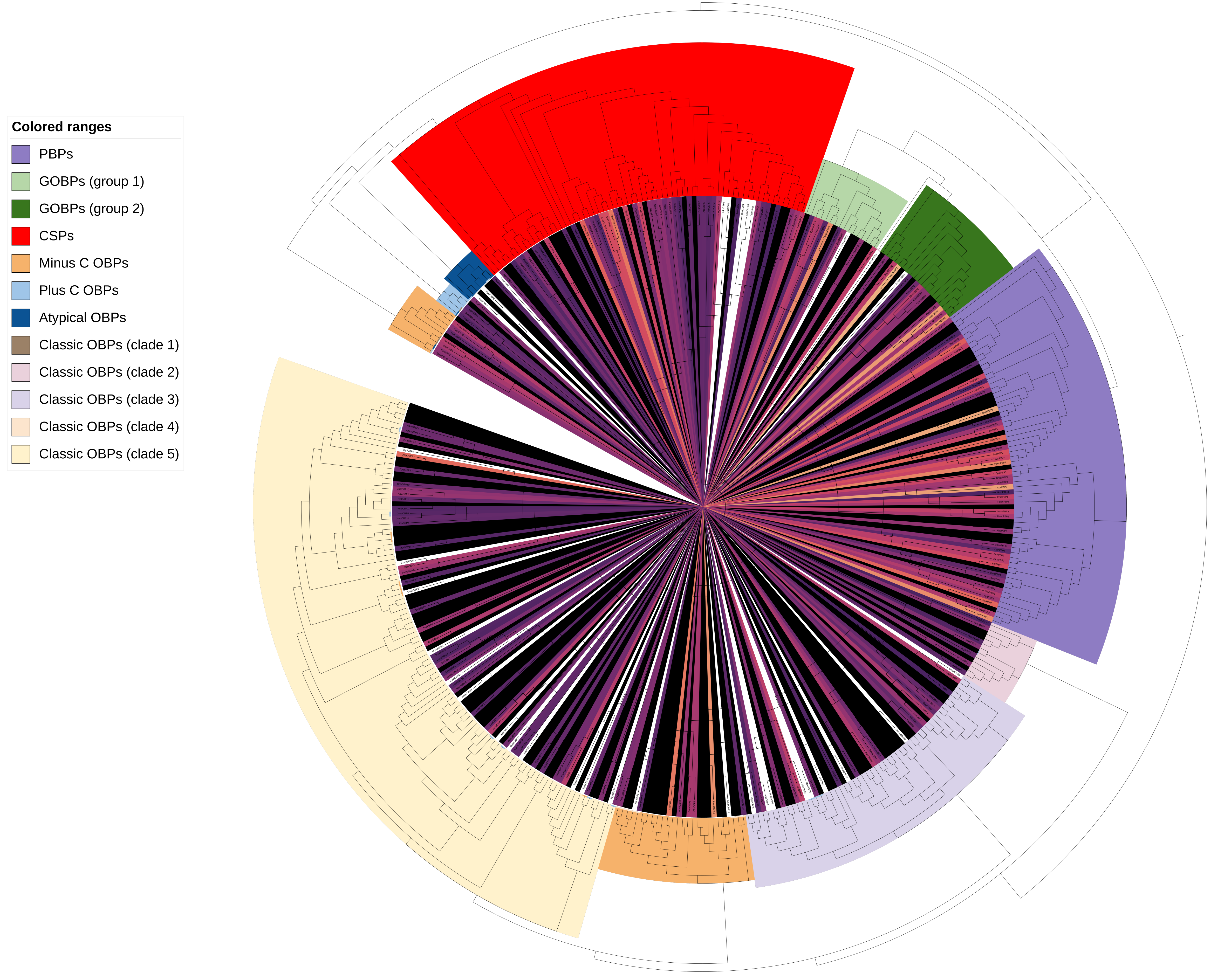

### Ester preference

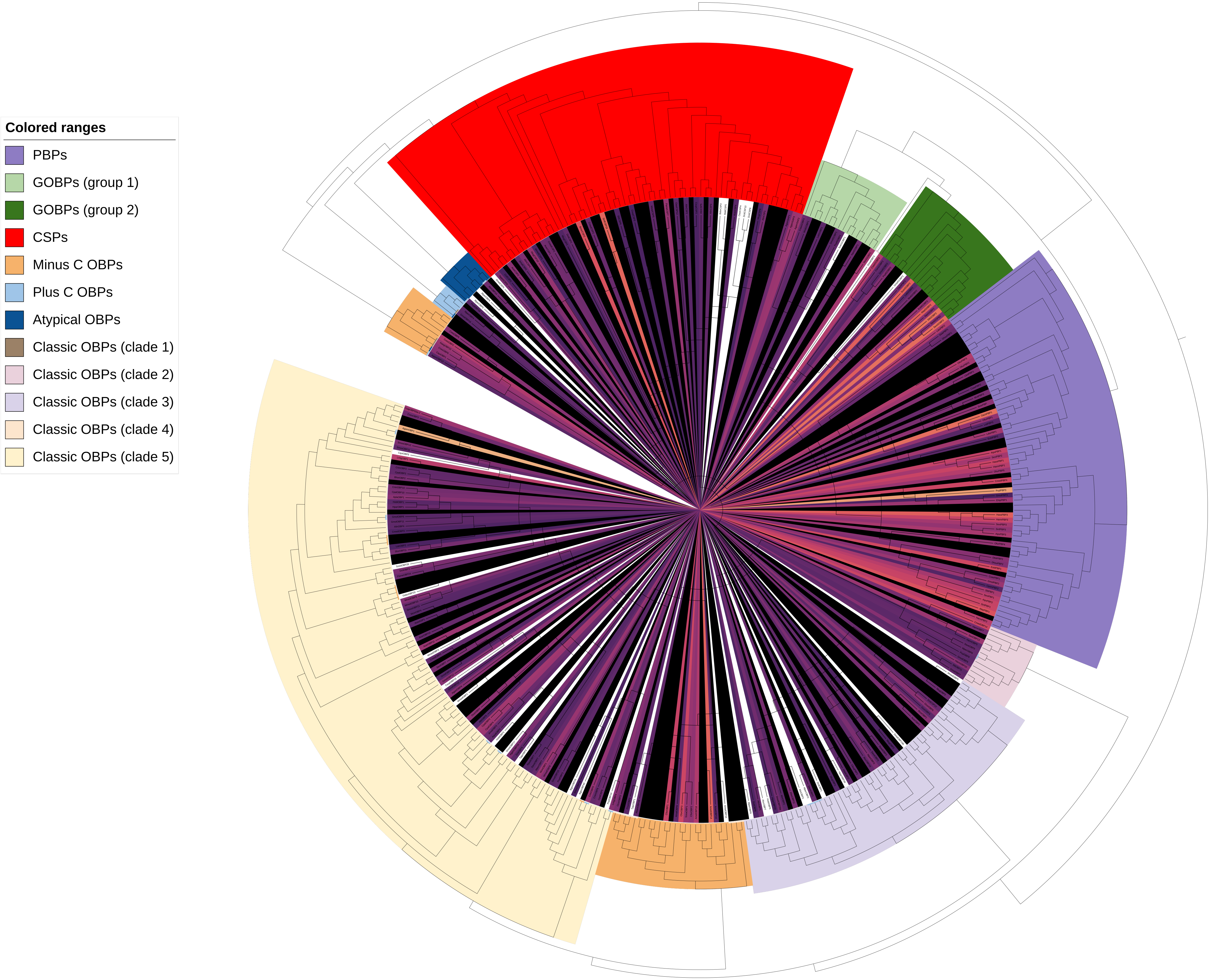

### Aromatic preference

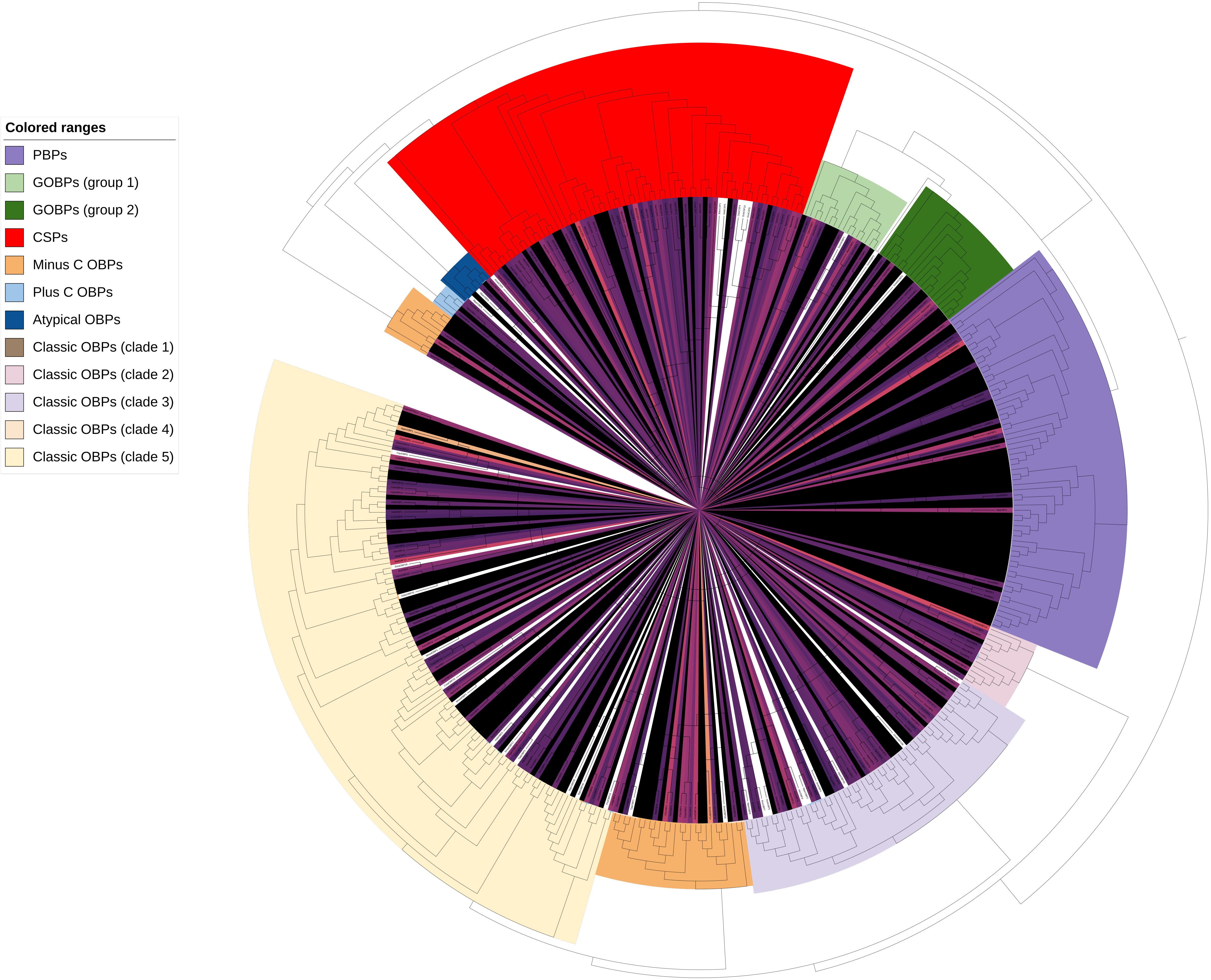
